# A fast and flexible approximation of power-law adaptation for auditory computational models

**DOI:** 10.1101/2023.11.30.569467

**Authors:** Daniel R. Guest, Laurel H. Carney

## Abstract

Power-law adaptation is a form of neural adaptation that has been shown to provide a better description of auditory-nerve adaptation dynamics as compared to simpler exponential-adaptation processes. However, the computational costs associated with power-law adaptation are high and, problematically, grow superlinearly with the number of samples in the simulation. This cost limits the applicability of power-law adaptation in simulations of responses to relatively long stimuli, such as speech, or in simulations for which high sampling rates are needed. Here, we present a simple approximation to power-law adaptation based on a parallel set of exponential-adaptation processes with different time constants, demonstrate that the approximation improves on an existing approximation provided in the literature, and provide updates to a popular phenomenological model of the auditory periphery that implements the new approximation.

## 2. Introduction

Computational models of the auditory system that allow users to simulate auditory responses to arbitrary sound-pressure waveforms are an increasingly important part of the auditory-science ecosystem. These models have proven invaluable for interrelating data across different species, modalities, and paradigms. For example, the phenomenological auditory-nerve (AN) model of Zilany, Bruce, and Carney (2014), hereafter the ZBC model, has been used extensively by several groups in recent years to pursue a wide range of psychophysical and physiological questions (Carney and McDonough, 2019; Bianchi et al., 2019; Maxwell et al., 2020; Saddler et al., 2021; Polonenko and Maddox, 2021; Zaar and Carney, 2022; Guest and Oxenham, 2022; Feather et al., 2022; Carney et al., 2023; Hamza et al., 2023; Lindboom et al., 2023; Brennan et al., 2023). Differences among models often reflect differences in the underlying goals of those who build them (Osses et al., 2022). Some models, such as the popular gammatone filterbank models, focus on speed, simplicity, and interpretability. Others, such as the ZBC model, instead focus on achieving a high level of detail and fidelity to physiological data. Such detail typically comes at a substantial computational cost, which can dissuade users from applying models to problems that require long or large-scale simulations. To ensure that physiologically realistic simulations remain within the realm of computational feasibility for relevant users, there is a continued need to find new strategies to improve model performance.

Recent versions of the ZBC model improved predictions of certain AN response features, such as synchrony to sinusoidal amplitude modulation (SAM) and the time course of post-offset response recovery, by modifying the model’s inner-hair-cell-AN synapse stage to include power-law adaptation (PLA) (Zilany et al., 2009). PLA is a form of neural adaptation that strikes a balance between adaptation with infinite memory of past responses, or perfect adaptation, and adaptation with a fixed time constant and exponential forgetting of past responses, or exponential adaptation (EA) (Drew and Abbott, 2006). Time-varying response rates ***r* (*t*)** are the rectified difference between an input ***s* (*t*)**and the output of an integrator ***I* (*t*):**

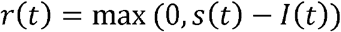

The integrator is governed by power-law temporal dynamics:

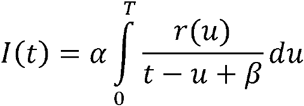

Although PLA improves the fidelity of the ZBC model to important aspects of AN physiology, its applicability is limited to short simulations at low underlying sampling rates because PLA is expensive to compute, and this expense grows superlinearly with the number of samples in the stimulus. In comparison, exponential adaptation can be implemented with a simple recursive relationship, wherein the output at each time step depends only on the current input and the output at the previous time step. In contrast, as can be seen in Equation 2, a direct implementation of PLA involves computing a weighted numerical integral over all previous time steps to determine the output value at each single time step. As a result, the computational costs of PLA grow exponentially with increasing simulation duration, whereas the computational cost of exponential adaptation is simply a constant proportion of stimulus duration. For very short simulations, the differences may be negligible; for simulations with long stimuli, or using very high sampling rates, a direct implementation of PLA becomes computationally prohibitive.

Previous versions of the ZBC model that included power-law adaptation avoided this problem in two ways. First, because the model’s IHC stage includes a cascade of lowpass filters that removes most energy beyond a few thousand Hz, the output of the IHC stage can be downsampled to 10 kHz before simulating the following stages of the model (i.e., IHC-AN synapse, AN spike generator) at a 10-kHz sampling rate. The outputs of the IHC-AN synapse and/or spike generator are then upsampled back to the original sampling rate before being returned to the user. This strategy significantly reduces the computational costs of PLA because only a fraction of output samples in the PLA adaptation stage are actually computed, while the rest are interpolated. Second, the ZBC model includes an approximation to PLA implemented as a two linear IIR filters (one filter for the “slow” PLA pathway and one for the “fast” PLA pathway in the model). Hereafter, this scheme is called “ZBC 2009/14 PLA”. The coefficients of the filters were numerically optimized to minimize differences between responses under true PLA and ZBC 2009/14 PLA (Zilany et al., 2009).

Here, we present an alternative scheme for approximating PLA. Instead of single, high-order IIR filters with coefficients that must be jointly optimized, we approximate PLA with a set of many parallel exponential-adaptation processes (Drew and Abbott, 2006). Hereafter, we refer to this scheme as “parallel-exponential PLA”. The idea is straightforward: to approximate the multi-time-scale behavior of PLA, one can use a finite number of exponential adaptation processes, each with a different time constant that mimics PLA dynamics at a particular time scale. Each exponential adaptation process can be easily implemented as a single-pole IIR lowpass filter. The time constants of each process are assumed to be distributed evenly in log-time between a short time constant (τ_*s*_)and a long time constant (τ_*l*_) resulting in just a few free parameters for each PLA pathway that can be selected manually or numerically optimized: τ_*s*_, (τ_*l*_), the number of separate processes (***N***), and a strength or scale parameter(***S***). We describe the implementation details for parallel-exponential PLA and show that it provides a fast and flexible approximation of PLA in the ZBC model.

## 3. Methods

### General simulation methods

Simulations of AN responses were conducted using the model of Zilany et al. (2014). Simulations used the parameter set intended to reproduce AN data from cats. Simulated fibers were always low-threshold high-spontaneous-rate fibers (Liberman, 1978). Except where otherwise specified, fractional Gaussian noise was included in the inner-hair-cell-AN synapse stage using default parameters for each model. PLA adaptation in the same synapse was implemented either using the native implementation provided in the model’s original source code (“true PLA”), the approximate implementation provided in the model’s original source code (“ZBC 2009/14 PLA”), or a new approximation based on a modified version of the source code described below (“parallel-exponential PLA”). Simulations were conducted with a sampling rate of 100 kHz.

### 3.1. Source-code modifications

Source code for the AN model of Zilany et al. (2014) was obtained online (<u>link</u>) and modified in the following ways. Parallel-exponential PLA was added to the synapse simulation code as an additional branch for implementing PLA, alongside true PLA and ZBC 2009/14 PLA. One free parameter in parallel-exponential PLA is the number of parallel exponential processes. For the present simulations, this number was fixed at 100. As before, time-varying response rate *r* [*n*] is the rectified difference between an input *s* [*n*]and the output of an integrator *I* [*n*]:

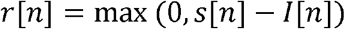

Here, *s* (*t*) is the output of the exponential-adaptation stage of the model (Zilany et al., 2009). The integrator is governed according to the equations:

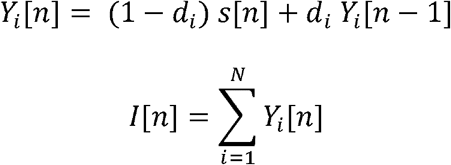

Where ***d***_*i*_ is the i-th decay coefficient for a single-pole lowpass IIR filter (determined by 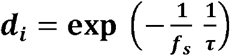, where **τ** is the time constant associated with the *i*-th process), and ***Y***_*i*_ is the state variable associated with the i-th process.

### 2.3. Numerical optimization

Parameter values for parallel-exponential PLA were selected by numerical optimization. First, waveforms were synthesized for a novel modulated stimulus, hereafter the “cycle-by-cycle roved sinusoidally amplitude-modulated (SAM) tone”, designed to challenge the ability of the approximate PLA schemes to accurately simulate the dynamics of PLA. The stimulus carrier was a pure tone. The stimulus modulator was a sinusoidal modulator with random periods of time drawn from a uniform distribution from 0 to 100 ms inserted as “deadtime” between each cycle of the modulator. After modulating the carrier with the modulator, each modulated period was identified and the overall sound pressure level of each modulated period was randomized over a uniform distribution from 20 to 70 dB SPL. The insertion of random level variations and random modulator gaps in what is otherwise a sinusoidally amplitude-modulated tone was intended to challenge the approximate PLA schemes’ ability to simulate the dynamics of PLA. Insertion of random modulator gaps produced onset-like responses at the start of each modulation period and also made post-offset recovery dynamics more salient. Insertion of random variations in sound level challenged the approximation to replicate how adaptation dynamics changed across the sound-level range. In both cases, the stimulus dynamics resulted in a substantial challenge for both approximations to model how PLA shaped both onset and offset responses as a function of stimulus duration, interstimulus interval, and sound level. The following parameter values were used during optimization: carrier frequencies of 1 and 10 kHz; modulation rates of 16 and 64 Hz; modulation depths of -10 and -5 dB; and durations of 1 and 5 s.

Next, inner-hair-cell responses were simulated for each stimulus waveform using the ZBC model. An 250 ms of additional time was simulated after stimulus offset to ensure that post-offset recovery was included as part of the simulations. The CF of the inner hair cell was set to match the carrier frequency of the stimulus. The inner-hair-cell waveforms were then used to simulate responses from the auditory-nerve stage of the ZBC model under true PLA. The same frozen sample of fractional Gaussian noise was used in the synapse model for all simulations to ensure that responses (and the resulting optimization procedure) were deterministic. The instantaneous-rate output was used instead of the spiking output for the same reason. High-spontaneous-rate fibers were simulated, but the estimated parameters could also be applied to simulations of medium- or low-spontaneous-rate fibers.

Parameters for parallel-exponential PLA were then optimized numerically by use of MATLAB’s *fmincon*. On each step of the procedure, current parameter-value estimates were used to simulate AN responses under parallel-exponential PLA, RMS error between the pre-computed responses with true PLA and responses with parallel-exponential PLA was computed for each stimulus waveform, and then RMS errors were summed across all stimulus waveforms to arrive at a final loss value.

## 4. Results

The results are reported in Fig. 1. For short duration simulations (< 500 ms), ZBC 2009/14 PLA and parallel-exponential PLA performed very similarly, both accurately reproducing the simulated dynamics of AN responses under PLA (Fig. 1a, left side). This result was expected, as the ZBC 2009/14 PLA approximation was designed for short simulations (Zilany et al., 2009). However, at longer time scales (> 2 s), substantial differences arose between true PLA and ZBC 2009/14 PLA: steady-state response magnitudes tended to be overestimated (Fig. 1a, right side, #1), and recovery from stimulation post-offset tended to be too fast, both after stimulation by low-level sounds, which tended to result in fast recovery (Fig. 1a, right side, #2), and after stimulation by high-level sounds, which tended to result in slower recovery (Fig. 1a, right side, #3). These trends are consistent with the IIR filters in the ZBC 2009/14 PLA being optimized to reproduce PLA behavior on short time scales, and underestimating the buildup of adaptation at longer time scales. In contrast, the parallel-exponential scheme remained accurate at these time scales, precisely reproducing the true-PLA response.

**Figure 1.**
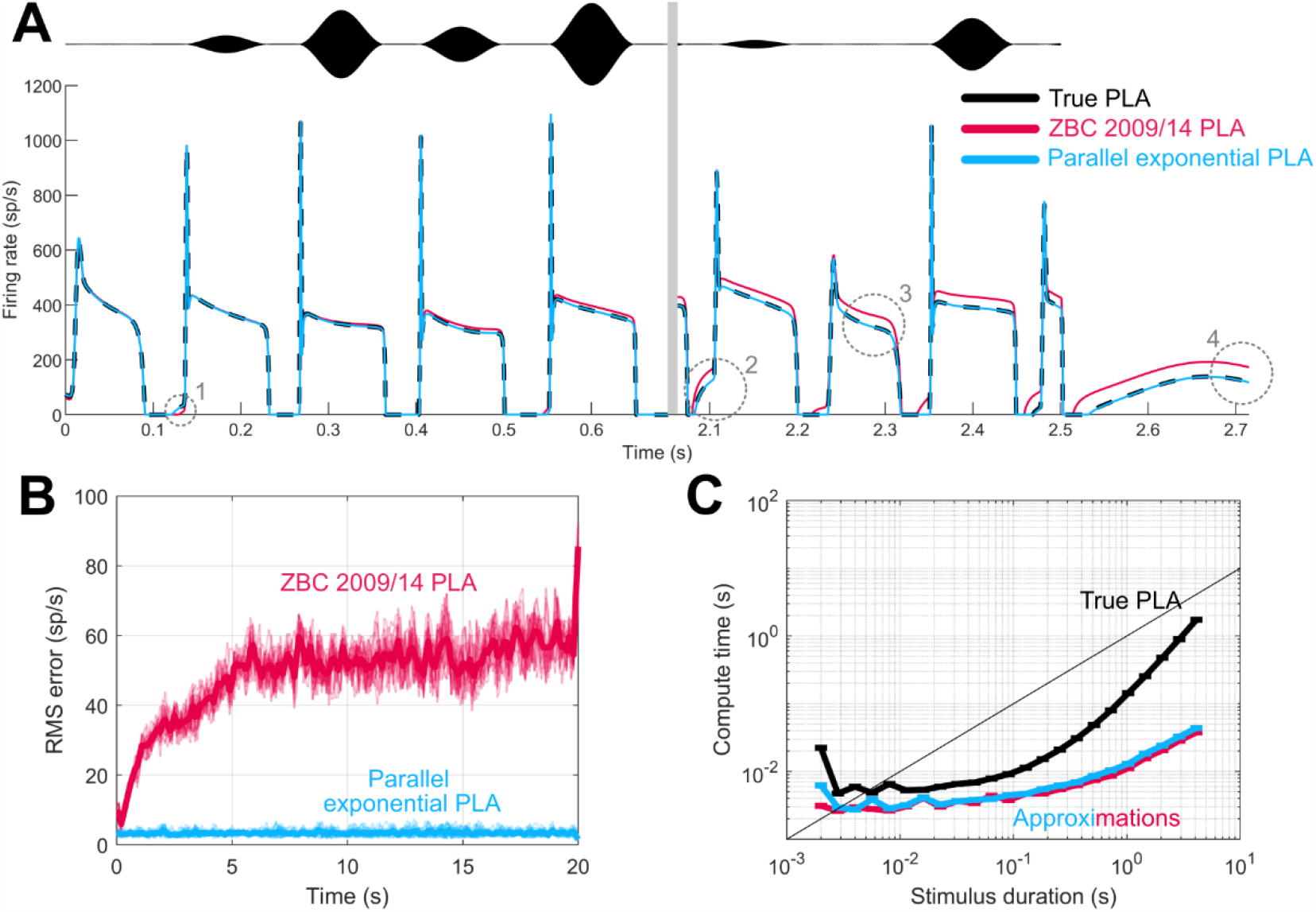
**A.** Instantaneous rate in response to a 2.5-s cycle-by-cycle roved SAM tone stimulus as a function of time for different implementations of PLA (color). The thin gray bar after 0.6 seconds indicates a time skip along the x-axis. The black trace at the top shows the stimulus waveform. Gray circles and numbers indicate important details referred to in the text. **B.** RMS error for approximate PLA in response to a 20-s cycle-by-cycle roved SAM tone as a function of time, windowed with a sliding 200-ms window, for different implementations of approximate PLA (color). Thin traces show results for single waveforms, thicker traces show the average over all tested waveforms. **C.** Compute time versus stimulus duration for different implementations of PLA (color). Color available online.

For much longer simulations (25 s) and simulations averaged over different samples of stimulus noise waveforms, parallel-exponential PLA produced approximations of true-PLA responses that retained low error across the entire time range (Fig. 1b), whereas ZBC 2009/14 PLA exhibited low-error only for the initial ∼1 s of the response. Importantly, despite improvements in fidelity, parallel-exponential PLA was about as expensive to compute as ZBC 2009/14 PLA (Fig. 1c). Both approximation schemes were substantially less expensive to compute than true PLA, especially for long simulations. For example, for a 100-ms simulation, simulating true PLA was only approximately 2.4 times slower than approximating PLA (runtime true PLA = 11.80 ms, runtime parallel exponential PLA = 4.82 ms). In contrast, for a 5-s simulation, simulating true PLA was dramatically more expensive, with a runtime approximately 37 times slower than approximating PLA (runtime true PLA = 2.85 s, runtime parallel exponential PLA 0.08 s).

It is clear from Fig. 1 that, in response to our novel stimulus, the new PLA approximation scheme provides a good qualitative and quantitative fit to model responses simulated using true PLA. However, it is still important to verify that the new scheme correctly emulates physiological responses that were used to fit the original model, including its PLA stage. One of the more important results of adding PLA to the model was an increase in modulation gain in response to SAM tones, quantified by measuring synchrony to the modulation frequency of a SAM tone as a function of modulation depth (Zilany et al., 2009). This increased modulation gain better matched AN physiological data, as compared to older versions of the model. As can be seen in Fig. 2, we repeated this simulation with true PLA, ZBC 2009/14 PLA, and parallel exponential PLA, and found that both approximations correctly exhibit the modulation-gain values of 5–10 dB seen under true PLA.

**Figure 2.**
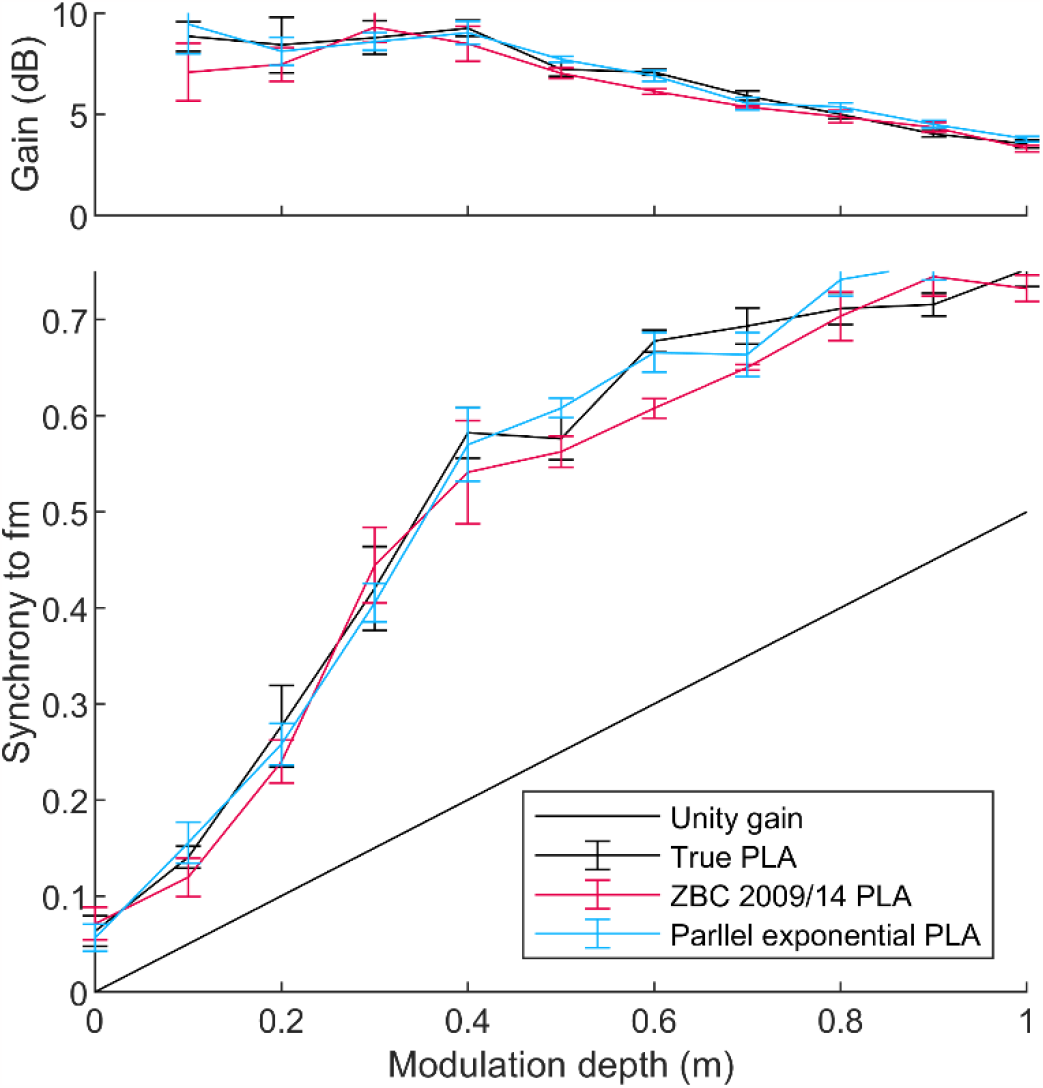
Top. Modulation gain as a function of modulation depth for each tested implementation of PLA in response to SAM tones with a 100-Hz modulation frequency presented at level of 22 dB SPL (approximately 5 dB above threshold) at a CF of 20.2 kHz, where modulation gain is quantified as **20 log**_**10**_**(2 *θ/m*)**, and ***θ*** is the vector strength at 100 Hz. **Bottom**. Vector strength for the same simulations. In both panels, results were simulated for five different repetitions. Means and ±1 standard error across those repetitions are plotted as points and error bars, respectively.

## 5. Discussion

PLA is an important aspect of AN adaptation, and here we provide a new scheme for approximating it that is efficient while preserving fidelity at long time scales. The primary applications for this approximation are simulations that involve long periods of time and simulations that require high sampling rates. The former is naturally encountered on a frequent basis in auditory simulations. For example, accurately simulating physiological protocols involves simulation of responses to both acoustic stimuli and interstimulus intervals between acoustic stimuli. That is, a model should receive a single continuous time-pressure waveform consisting of alternating acoustic stimulation and gaps of silence, to account for the persistence of adaptation across time and recovery from adaptation in between stimuli. Such simulations are prohibitively slow using true PLA, but parallel-exponential PLA is well suited to this usage. The latter is often encountered when working with high acoustic frequencies, when working with ideal-observer models that can be sensitive to the effects of aliasing (e.g., Heinz et al., 2001), or when simulating dynamic feedback loops that preclude intermediate downsampling steps to increase efficiency (Farhadi et al., 2023). Future work could explore better ways to fit the PLA approximation parameters and the optimal number of discrete processes to use in the approximation scheme to balance between computational costs and fidelity to true PLA.

## Acknowledgments

This work was supported by NIH-NIDCD-R01010813.

## Data Availability

Necessary code for reproducing this manuscript is available online at [[link forthcoming]].

## References

Bianchi, F., Carney, L. H., Dau, T., and Santurette, S. (2019). “Effects of Musical Training and Hearing Loss on Fundamental Frequency Discrimination and Temporal Fine Structure Processing: Psychophysics and Modeling,” Journal of the Association for Research in Otolaryngology. doi:10.1007/s10162-018-00710-2

Brennan, M. A., Svec, A., Farhadi, A., Maxwell, B. N., and Carney, L. H. (2023). “Inherent envelope fluctuations in forward masking: Effects of age and hearing loss,” The Journal of the Acoustical Society of America, 153, 1994. doi:10.1121/10.0017724

Carney, L. H., Cameron, D. A., Kinast, K. B., Feld, C. E., Schwarz, D. M., Leong, U.-C., and McDonough, J. M. (2023). “Effects of sensorineural hearing loss on formant-frequency discrimination: Measurements and models,” Hearing Research, 435, 108788. doi:10.1016/j.heares.2023.108788

Carney, L. H., and McDonough, J. M. (2019). “Nonlinear auditory models yield new insights into representations of vowels,” Attention, Perception, & Psychophysics, 81, 1034–1046. doi:10.3758/s13414-018-01644-w

Drew, P. J., and Abbott, L. F. (2006). “Models and Properties of Power-Law Adaptation in Neural Systems,” Journal of Neurophysiology, 96, 826–833. doi:10.1152/jn.00134.2006

Farhadi, A., Jennings, S. G., Strickland, E. A., and Carney, L. H. (2023). “Subcortical auditory model including efferent dynamic gain control with inputs from cochlear nucleus and inferior colliculus,” The Journal of the Acoustical Society of America. doi:10.1121/10.0022578

Guest, D. R., and Oxenham, A. J. (2022). “Human Discrimination and Modeling of High-Frequency Complex Tones Shed Light on the Neural Codes for Pitch,” PLOS Computational Biology, 18, e1009889. doi:10.1371/jounral.pcbi.1009889

Heinz, M. G., Colburn, H. S., and Carney, L. H. (2001). “Evaluating Auditory Performance Limits: I. One-Parameter Discrimination Using a Computational Model for the Auditory Nerve,” Neural Computation, 13, 2273–2316. doi:10.1162/089976601750541804

Liberman, M. C. (1978). “Auditory□nerve Response From Cats Raised in a Low□noise Chamber,” The Journal of the Acoustical Society of America, 63, 442–455. doi:10.1121/1.381736

Lindboom, E., Nidiffer, A., Carney, L. H., and Lalor, E. C. (2023). “Incorporating models of subcortical processing improves the ability to predict EEG responses to natural speech,” Hearing Research, 433, 108767. doi:10.1016/j.heares.2023.108767

Maxwell, B. N., Richards, V. M., and Carney, L. H. (2020). “Neural fluctuation cues for simultaneous notched-noise masking and profile-analysis tasks: insights from model midbrain responses,” The Journal of the Acoustical Society of America, 147, 3523–3537. doi:10.1121/10.0001226

Polonenko, M. J., and Maddox, R. K. (2021). “Exposing distinct subcortical components of the auditory brainstem response evoked by continuous naturalistic speech,” eLife, doi: 10.7554/elife.62329. doi:10.7554/elife.62329

Saddler, M. R., Gonzalez, R., and McDermott, J. H. (2021). “Deep Neural Network Models Reveal Interplay of Peripheral Coding and Stimulus Statistics in Pitch Perception,” Nature Communications, 12, 7278. doi:10.1038/s41467-021-27366-6

Vecchi, A. O., Varnet, L., Carney, L. H., Dau, T., Bruce, I. C., Verhulst, S., and Majdak, P. (2021). “A Comparative Study of Eight Human Auditory Models of Monaural Processing,” arXiv.

Zaar, J., and Carney, L. H. (2022). “Predicting speech intelligibility in hearing-impaired listeners using a physiologically inspired auditory model,” Hearing Research, 426, 108553. doi:10.1016/j.heares.2022.108553

Zilany, M. S. A., Bruce, I. C., and Carney, L. H. (2014). “Updated Parameters and Expanded Simulation Options for a Model of the Auditory Periphery,” The Journal of the Acoustical Society of America, 135, 283–286. doi:10.1121/1.4837815

